# MtrAB two-component system is crucial for the intrinsic resistance and virulence of *Mycobacterium abscessus*

**DOI:** 10.1101/2024.04.05.588258

**Authors:** Jingran Zhang, Yanan Ju, Lijie Li, Adnan Hameed, Buhari Yusuf, Yamin Gao, Cuiting Fang, Xirong Tian, Jie Ding, Wanli Ma, Xinwen Chen, Shuai Wang, Tianyu Zhang

## Abstract

*Mycobacterium abscessus* (Mab) poses serious therapeutic challenges, principally due to its intrinsic resistance to a wide array of antibiotics. The pressing issue of drug resistance has spurred an urgent need to explore novel targets and develop new therapeutic agents against Mab. The MtrAB two-component system, conserved among Actinobacteria, is pivotal for regulating various metabolic processes. Nevertheless, the role of MtrAB in Mab remains elusive.

In this study, we uncovered that Mab strains with disrupted *mtrA, mtrB* or both exhibited heightened susceptibility to a variety of antibiotics with diverse mechanisms of action, in contrast to the wild-type strain. In a murine model, rifabutin, bedaquiline, and amikacin, which were inactive against the wild-type Mab strain, demonstrated efficacy against all the *mtrA, mtrB* and *mtrAB* knockout strains, significantly reducing pulmonary bacterial burdens compared to vehicle controls after ten days of treatment. Notably, the virulence of all the *mtrA, mtrB*, and *mtrAB* knockout strains was highly diminished in the murine model, as evidenced by a substantial decrease in bacterial load in the lungs of mice after 16 days. We observed that all three knockout strains exhibited a significantly reduced growth rate compared to the wild-type strain.

We discovered that cells lacking *mtrA, mtrB* or both exhibited an elongated cell length and had multiple septa, suggesting that both MtrA and MtrB regulate cell division of Mab. Subsequently, an ethidium bromide accumulation assay disclosed that the absence of either *mtrA* or *mtrB* or both significantly increased cell envelope permeability.

In summary, this study suggests that *mtrA* and *mtrB* play an important role in the intrinsic resistance and virulence of Mab by affecting cell division and altering cell permeability. Consequently, MtrA and MtrB represent promising targets for the discovery of anti-Mab drugs.

**HIGHLIGHTS:**

- Knockout of *mtrA, mtrB* or *mtrAB* leads to increased sensitivity of *M. abscessus in vitro* and *in vivo*.
- The *mtrA, mtrB* or *mtrAB* knockout *M. abscessus* strains exhibit highly reduced virulence.
- MtrA and MtrB are potential targets for anti-*M. abscessus* drug discovery.
- Knockout of *mtrA, mtrB* or *mtrAB* results in defective cell division in *M. abscessus*.

## 1. Introduction

*Mycobacterium abscessus* (Mab) is an emerging opportunistic pathogen causing skin, soft tissue and pulmonary infections in immunocompromised individuals and patients with pre-existing lung diseases such as cystic fibrosis and chronic obstructive pulmonary diseases[1]. This bacterium represents the predominant species among rapidly growing mycobacteria isolated from the lungs of cystic fibrosis patient[2–4]. The management of Mab infections poses substantial challenges, earning it the reputation of an “incurable nightmare”[5, 6], with current treatment regimens— comprising a multidrug regimen of macrolides, amikacin (AMK), cefoxitin (CFX), and imipenem[7] over an 18-24 months period—achieving a cure rate of merely ∼58%[8, 9]. The main reason for this dilemma is the high-level intrinsic resistance of Mab to nearly all antibiotics used in clinical practice, particularly including almost all antituberculosis drugs. Intrinsic resistance of Mab is mediated by multiple factors, including the multi-drug efflux pumps, numerous enzymes that can modify either the drug target or the drug itself, and the highly impermeable cell envelope[10]. Hence, studies of novel targets of Mab to identify effective combinations should be further emphasized.

Two-component system (TCS) is a regulatory mechanism commonly utilized by prokaryotes to regulate the expression of relevant genes in response to a variety of environmental stressors. A prototypical TCS comprises a histidine kinase (HK) that detects signal stimuli and a corresponding response regulator (RR) that can phosphorylated by activated HK. This activated RR then binds to specific DNA motifs within the promoter region, leading to alterations in gene expression[11]. Bacterial TCS is involved in regulating various cellular life processes, encompassing growth, virulence, persistence, and drug resistance. Additionally, due to their relatively conserved kinase domains, they represent an appealing target for the development of novel therapeutic agents. In *Mycobacterium tuberculosis* (Mtb), the DosRST system, which distinctively contains two HKs rather than the customary one, governs the conversion between dormancy and active state[12]. Zheng et al. recently discovered three novel inhibitors of the DosRST regulon via a high-throughput screening of approximately 540,000 small-molecule compounds library. Under hypoxia, all three compounds were found to inhibit physiological processes associated with Mtb persistence, including triacylglycerol synthesis, survival and antibiotic tolerance[13].

The MtrAB TCS, comprising the response regulator MtrA and histidine kinase MtrB, is conserved across Actinobacteria, including species such as *Dietzia* species, *Streptomyces, Mycobacterium* and *Corynebacterium glutamicum*[14–17]. The MtrAB TCS plays diverse roles in bacterial life activities. Specifically, in *Dietzia* sp. DQ12-45-1b, it regulates the cell wall homeostasis responding to environmental alkaline stress by modulating the expression of regulator MraZ[17]. In *Corynebacterium glutamicum,* the MtrAB TCS is involved in osmoregulation and cell wall metabolism[14]. Moreover, in *Streptomyces coelicolor*, it plays an important role in the developmental life cycle[16]. Additionally, in Mtb, MtrAB is an essential TCS involved in cell replication and division[18, 19]. Furthermore, MtrAB has been reported to be involved in drug resistance. The absence of *mtrAB* in *Corynebacterium glutamicum* leads to heightened susceptibility to penicillin and vancomycin (VAN), a phenomenon analogous to the increased sensitivity observed upon the deletion of *mtrB* in *Mycobacterium avium*[14, 20]. Comparably, in both Mtb and *Mycobacterium smegmatis* (Msm), the knockdown and knockout of *mtrA,* respectively, resulted in altered sensitivity to various drugs[18, 21].

However, in-depth studies of Mab TCS, particularly the MtrAB, have not been performed. Although the homology of MtrA in Mab and Mtb is high (91.23% identity) (Figure. S1), it is nonessential for the survival of Mab, contrary to its essential role in Mtb. In addition, given the diverse conditions that Mab encounters as an environmental opportunistic pathogen, coupled with its larger genome size and content compared to Mtb[22], it is highly plausible that the *mtrAB* cluster encompasses a unique set of genes implicated in various cellular processes.

In this study, we explored the functions of MtrA and MtrB within Mab, emphasizing their impact on intrinsic resistance and virulence. Moreover, we evaluated their potential as drug targets through *in vitro* and *in vivo* studies. This research endeavors to appraise their therapeutic potential, offering insights that may aid in the discovery and advancement of novel antimicrobial agents against this challenging pathogen.

## 2. Materials and Methods

### 2.1 Bacterial strains and culture condition

Mab *subsp. abscessus* GZ002 (NCBI GenBank accession number CP034181), previously described clinical isolate[23], was cultured at 37℃ in 7H9 medium supplemented with 0.2% glycerol, 0.05% Tween 80 and 10% oleic acid albumin dextrose catalase (OADC, Difco)) or 7H10 containing 0.2% glycerol and 10% OADC. When necessary, selection antibiotics kanamycin and zeocin were added to the medium at final concentrations of 100 μg/mL and 30 μg/mL respectively. The inducer anhydrotetracycline (ATc) was used at a concentration of 100 ng/mL. *Escherichia coli* DH5α was cultured at 37℃ in Luria Bertani (LB) broth or on LB agar. The antibiotic concentrations used for *Escherichia coli* were kanamycin 50 μg/mL, zeocin 30 μg/mL and ampicillin 100 μg/mL. The strains used in this study are listed in Table. S1.

### 2.2 Mab transposon (Tn) mutants screening and identification of Tn insertion sites

In reference to the previously published articles[24], Tn mutants were replica-plated on 7H10 agar, both with and without a subinhibitory concentration of RIF (4 μg/mL). Colonies exhibiting defective growth in the presence of RIF were harvested for drug susceptibility testing. The Tn insertion sites in mutants were determined as described previously[24]. Briefly, genomic DNA was extracted from a mixture of 10 Tn mutants, followed by next-generation sequencing (NGS) to locate Tn insertion sites within the genomes. Reads containing the transposon sequence were mapped to the genome, and only those insertions with read counts exceeding 20 at specific sites were deemed authentic. To corroborate the NGS findings, the disrupted region was amplified using PCR, employing a mixture of several colonies as a template in a single reaction, facilitated by primers that targeted both the Tn and the disrupted gene.

### 2.3 Deletion and complementation of *MAB_3591c, MAB_3590c, MAB_3590c*- *MAB_3591c* in Mab

Deletion of *MAB_3591c* was achieved using recombineering and the Xer/dif system as previously described[24, 25]. Briefly, the upstream and downstream flanking arms of *MAB_3591c* were amplified and then cloned on either side of the zeocin resistance gene within the vector pBluescript II SK(+). As shown in Figure. S2, the allelic exchange substrate, which includes the upstream and downstream regions along with the selection maker, was PCR-amplified from the vector backbone and electroporated into freshly prepared electrocompetent Mab cells carrying pJV53 plasmid. After a 5-day incubation period at 37℃, transformants that grew on 7H10 plates containing kanamycin and zeocin were identified via PCR. The correct *MAB_3591c* deletion colony was cultured in 7H9 medium without antibiotics for two generations to facilitate the loss of free plasmid pJV53 and the clearance of the zeocin resistance gene by the Xer/dif system. The marker-free deletion colony, named ΔmtrA, were identified by PCR and subsequently verified by sequencing (Figure. S2).

The complementation of the *MAB_3591c* gene was achieved by introducing pMV261-based plasmids expressing *MAB_3591c* into ΔmtrA, resulting in the strain CΔmtrA.

Deletion of *MAB_3590c* or *MAB_3590c-MAB_3591c* was carried out through CRISPR-associated recombineering as described earlier[26, 27]. Briefly, two complementary oligonucleotides containing the target sequence in *MAB_3590c* adjacent to PAM sequences YTN (position 904-906) were synthesized, annealed to yield a protospacer cassette with *Bpm* I and *Hin*d III overhangs at the 5′ and 3′ ends, respectively, and then cloned into pCR-Zeo. To obtain homologous arms, the vector pBluescript II SK(+) only carrying up and down homologous arms of the target gene (*MAB_3590c* or *MAB_3590c-MAB_3591c)* but lacking zeocin resistance gene was constructed. The allelic exchange substrate and crRNA-expressing pCR-Zeo plasmid were simultaneously electroporated into Mab cells harboring pJV53-Cpf1 plasmid. After 5 days of incubation at 30℃, transformants growing on 7H11 plates supplemented with kanamycin, zeocin and ATc were identified by PCR and confirmed by sequencing (Figure. S2).

Complementation of the *mtrB* and *mtrAB* knockout strains was achieved by expressing a single gene (*mtrA* or *mtrB*) or both genes using the pMV261 plasmid. All PCR primers and plasmids utilized in this study are detailed in Table. S3.

### 2.4 Drug susceptibility testing

The minimum inhibitory concentrations (MICs) were determined by the microdilution method in 96-well plates using Middlebrook 7H9 medium, without Tween 80. Bacteria were cultured to mid-log phase and subsequently diluted to achieve a density of 5 × 10^5^ colony forming units (CFUs)/mLin 7H9 medium containing two-fold serial dilutions of the drugs. The cultures were then incubated at 37℃ for three days. The MIC was defined as the lowest antibiotic concentration capable of inhibiting visible bacterial growth. The drugs utilized in this study for the *in vitro* experiments, such as linezolid (LZD), rifabutin (RFB), bedaquiline (BDQ), rifampicin (RIF), and clofazimine (CLF), were dissolved in dimethyl sulfoxide (DMSO). The concentration of the stock solution was determined based on maintaining a final DMSO concentration typically between 0.1% and 1% during culture. Meanwhile, levofloxacin (LEV), moxifloxacin (MXF), AMK, CFX, and VAN were solubilized in water and filtered through a 0.22 μm filter.

For the plate assay, Mab strains were cultivated to exponential phase, and 1 μL of tenfold serial dilutions was respectively dispensed onto 7H10 plates containing various drugs or no drugs. Each experiment was performed in triplicate and repeated three times.

### 2.5 Mouse infection and treatment

Female BALB/c mice, aged 6-8 weeks and weighing 19-22 grams (GemPharmatech), were used in this study. For effective immunosuppression, all mice received daily doses of dexamethasone (DEXA) for one week before infection and throughout the experiment. DEXA (D1756, Sigma-Aldrich) was dissolved in sterile phosphate-buffered saline (PBS) and administered subcutaneously at a dosage of 4 mg/kg/day, as described previously[28]. Mice were infected via inhalation with wild-type Mab, *mtrA, mtrB*, or both *mtrA* and *mtrB* knockout strains, respectively, at the logarithmic growth stage. Three hours post-infection, four mice from each group were euthanized to determine the initial bacterial load in the lungs. The organs were homogenized in PBS, and serial 10-fold dilutions were plated onto 7H10 agar for CFU enumeration.

### 2.6 *In vivo* growth rate measurement

On days 6, 11 and 16 after infection, every four mice infected with different strains without any treatment except the DEXA, were euthanized, and lungs were removed aseptically for CFU enumeration.

### 2.7 Drug regimens

The mice were randomly assigned to receive different treatments starting from the day after infection. These treatments included daily oral gavage treatments of LZD at 100 mg/kg, RFB at 10 mg/kg, BDQ at 20 mg/kg, and MXF at 100 mg/kg, as well as subcutaneous injections of AMK at 150 mg/kg. LZD and RFB were dissolved in 0.4% carboxymethylcellulose sodium salt (CMC-Na), while BDQ was prepared in a solution of 20% hydroxypropyl-β-cyclodextrin solution acidified with 1.5% 1 mol/L HCl. MXF and AMK were dissolved in sterile water and PBS respectively. After 10 days of treatment, the mice were sacrificed for CFU determination in their lungs. The 7H10 agar for the BDQ treatment group was prepared with additional activated carbon (0.4%, wt/vol) for avoiding the carry over.

### 2.7 *In vitro* growth rate measurement

Bacterial growth was monitored by measuring changes in absorbance at 600 nm, and viability was determined by assaying CFU/mL. Mab cultures were grown to exponential phase and diluted to OD_600_ = 0.05, then incubated at 37℃ in a shaker at 200 rpm. OD_600_ was measured every 6 hours for 78 hours using a multifunctional microplate reader system (Molecular Device, FlexStation3). CFU were calculated by diluting the bacteria at indicated times, spreading the diluent onto a 7H10 agar plate and counting after 5-7 days of incubation at 37℃.

### 2.8 Microscopy and image analysis

The exponential phase bacterial cells were washed three times with 1 × PBS buffer (pH 7.2) to remove the medium and collected by centrifugation at 5,000 rpm for 5 min at 4℃. The bacterial cells were fixed 1 h at room temperature using 4% glutaraldehyde. Afterwards, glutaraldehyde was discarded, and cells were rinsed three times with PBS and dehydrated through a graded series of ethanol (70, 80, 90 and 100%) followed by critical point drying with liquid CO_2_. The dried specimens were then sputter-coated with aurum and imaged via Cryo-scanning electron microscopy (GeminiSEM). Cell lengths were analyzed using the Nano Measure software.

### 2.9 Cell envelope permeability assay

Cell envelope permeability of various Mab strains was assessed using the ethidium bromide (EtBr) uptake assay as previously described[24]. In brief, cultures of Mab in the mid-log phase were washed once in PBS containing 0.05% Tween 80 and adjusted to an OD_600_ of 0.8 in PBS supplemented with 0.8% glucose. A volume of 100 μL of bacteria was added to triplicate wells in a light-proof 96-well plate, and an equal volume of PBS with 4 μg/mL EtBr was added to each well, resulting in a final concentration of 2 μg/mL EtBr and 0.4% glucose. The fluorescence of EtBr was measured using a multifunctional microplate reader system (Molecular Device, FlexStation3) at 1-min intervals over a course of 60 min, with excitation and emission wavelengths set at 530 and 590 nm, respectively.

### 2.10 Comparison of bacterial sedimentation rates

Mab strains are cultured until they all attain the same log phase of growth. An equivalent volume of the bacterial liquid from each strain is then drawn and placed into separate centrifuge tubes, with photographic documentation capturing the initial appearance of the cultures. After a static incubation at room temperature for four hours, photographs are taken once more to observe and record the sedimentation of the bacterial cells.

## 3. Results

### 3.1 I5 Tn Mab mutant demonstrated an increased susceptibility to a variety of drugs

From our previously constructed Mab Tn mutant library, we screened 10 mutants exhibiting hypersensitivity to RIF, one of which was named as I5[24]. The microdilution method confirmed the hypersensitive phenotype, revealing that the MIC of RIF against the I5 strain was 1/16th of the wild-type Mab’s. Subsequently, we assessed the susceptibility of I5 to a range of antibiotics with distinct mechanisms of action. Notably, it demonstrated increased susceptibility to several drugs, including translation inhibitors such as LZD, RFB, RIF and AMK; cell envelope synthesis inhibitors like VAN and CFX; fluoroquinolone drugs affecting DNA replication, namely LEV and MXF; and respiratory chain targeting drugs CLF and BDQ (Figure. 1 and Table. 1).

**Figure 1.**
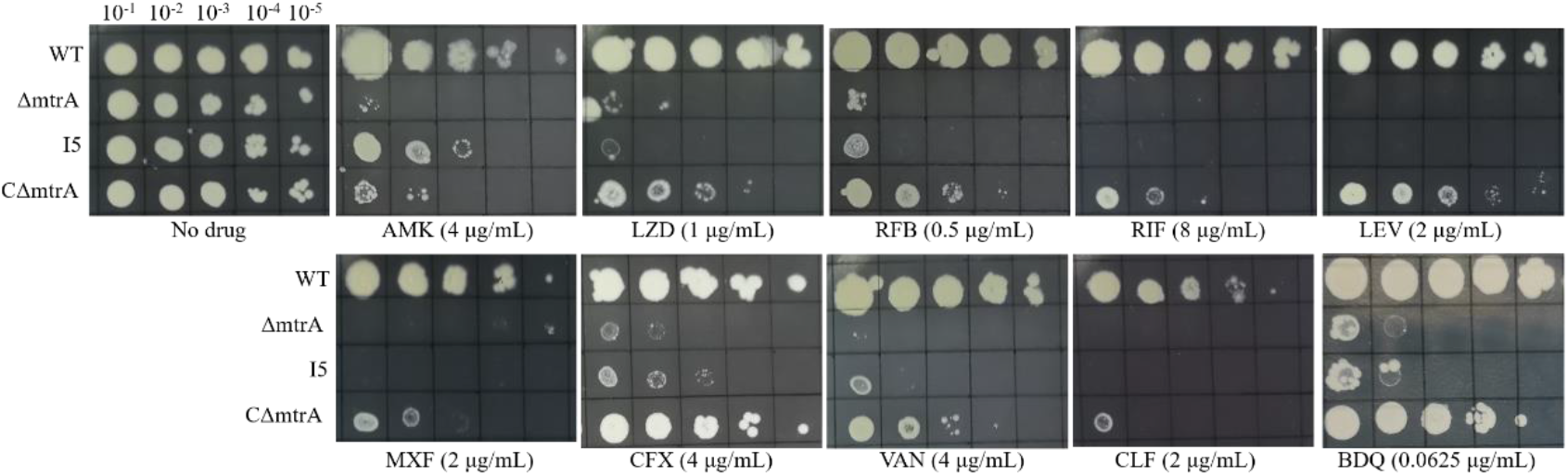
The disruption of *mtrA (MAB_3591c)* leads to heightened susceptibility to a variety of drugs. The abbreviations used are as follows: WT refers to wild-type Mab, ΔmtrA is the *mtrA* knockout Mab strain, I5 represents the transposon mutant Mab strain, and CΔmtrA is the complementary Mab strain, ΔmtrA carrying pMV261-mtrA expressing *mtrA* under the strong mycobacterial promoter *hsp60*. One microliter of 10-fold dilution series serially diluted bacterial cultures of exponentially growing cultures was spotted on Middlebrook 7H10 agar plates in the absence or presence of drugs at the indicated concentrations. The plates were then incubated at 37℃ for 5 days. The data presented are representative of the outcomes from three separate independent experiments.

**Table 1.** MICs of various drugs against different Mab strains.

| Drugs | Mab strains/ <sup>a</sup>MICs ($\mu\text{g/mL}$) | | | |
| --- | --- | --- | --- | --- |
| | WT | $\Delta mtrA$ | I5 | C$\Delta mtrA$ |
| Linezolid | 64 | 0.5-1 | 0.5-1 | 16 |
| Amikacin | 16 | 2-4 | 4 | 16 |
| Cefoxitin | 32 | 2-4 | 4 | 32 |
| Clofazimine | 4 | 0.5 | 1 | 2 |
| Levofloxacin | 32 | 2-4 | 4 | 8 |
| Moxifloxacin | 8-16 | 2-4 | 4 | 8 |
| Rifabutin | 4 | 1 | 1 | 2 |
| Rifampin | >128 | 8 | 4 | 32 |
| Bedaquiline | 0.5 | 0.0625 | 0.0625 | 0.25 |
| Vancomycin | 32 | 2-4 | 2 | 8-16 |

<sup>a</sup>MIC was defined as the lowest antibiotic concentration capable of inhibiting visible bacterial growth. WT, wild-type Mab; ΔmtrA, the *mtrA* knockout Mab; I5, the transposon mutant; CΔmtrA, the complementary strain of ΔmtrA carrying pMV261-*mtrA* expressing *mtrA* under the strong mycobacterial promoter *hsp60*. The experiment was executed in triplicate and repeated three times.

We identified the Tn-disrupted gene in the I5 strain as the *EFV83_04085* gene, which corresponds to *MAB_3591c* in Mab ATCC 19977. The Tn inserted into the “TA” dinucleotide located, 282 bases from the start codon of the 678-bp-long *MAB_3591c*.

### 3.2 Deletion of *MAB_3591c* resulted in hypersensitivity of Mab to multiple antibiotics *in vitro*

The amino acid sequence of MAB_3591 shares 93.86% identity to Mtb MtrA and 91.23% identity to Msm MtrA (Figure. S1), suggesting that MAB_3591 likely functions as MtrA in Mab. To verify that the increased sensitivity of I5 resulted from the disruption of *MAB_3591c*, a selectable marker-free *MAB_3591c* in-frame knockout strain (ΔmtrA) was further constructed using recombineering and the Xer/dif system as previously described[25, 29]. The correct colony was verified by PCR and sequencing (Figure. S2). Similarly to the Tn-mutant I5, ΔmtrA exhibited a heightened susceptibility to AMK, LZD, RFB, RIF, LEV, MXF, CFX, VAN, CLF, and BDQ when compared to the wild-type Mab (Figure. 1 and Table. 1). Complementation of *mtrA* (CΔmtrA) restored the antibiotic resistance, indicating that *mtrA* plays a critical role in establishing resistance to antibiotics stress in Mab (Figure. 1 and Table. 1).

### 3.3 Deletion of *mtrB* and *mtrAB* hypersensitizes Mab to multiple antibiotics *in vitro*

MtrA, potentially functioning as a response regulator within the MtrAB TCS, requires phosphorylation and activation by MtrB for its initial functional step. To elucidate the role of MtrAB, we constructed the *mtrB* (*MAB_3590c*) knockout strain, denoted as ΔmtrB, and the *mtrAB* (*MAB_3590c-MAB_3591c*) double knockout strain, referred to as ΔmtrAB further, utilizing CRISPR-associated recombineering techniques[25, 26]. We then comparatively evaluated the *in vitro* drug sensitivity of these knockout strains. The deletion of *mtrB* resulted in heightened drug sensitivity surpassing that of wild-type (WT), akin to ΔmtrA (Figure. 2A and Table. 2). The complementation of *mtrB* (CΔmtrB) restored the antibiotic resistance. Furthermore, ΔmtrAB exhibited an augmented sensitivity beyond that of the individual knockout strains ΔmtrA and ΔmtrB (Figure. 2B and Table. 3). Upon the combined complementation of *mtrA* and *mtrB* (CΔmtrAB), drug resistance was effectively restored. These findings indicate that the MtrAB TCS plays a role in modulating the intrinsic resistance of Mab. Notably, overexpression of mtrA in ΔmtrB or ΔmtrAB backgrounds resulted in the restoration of drug resistance, as shown in Figure 2B and Table 3.

**Figure 2.**
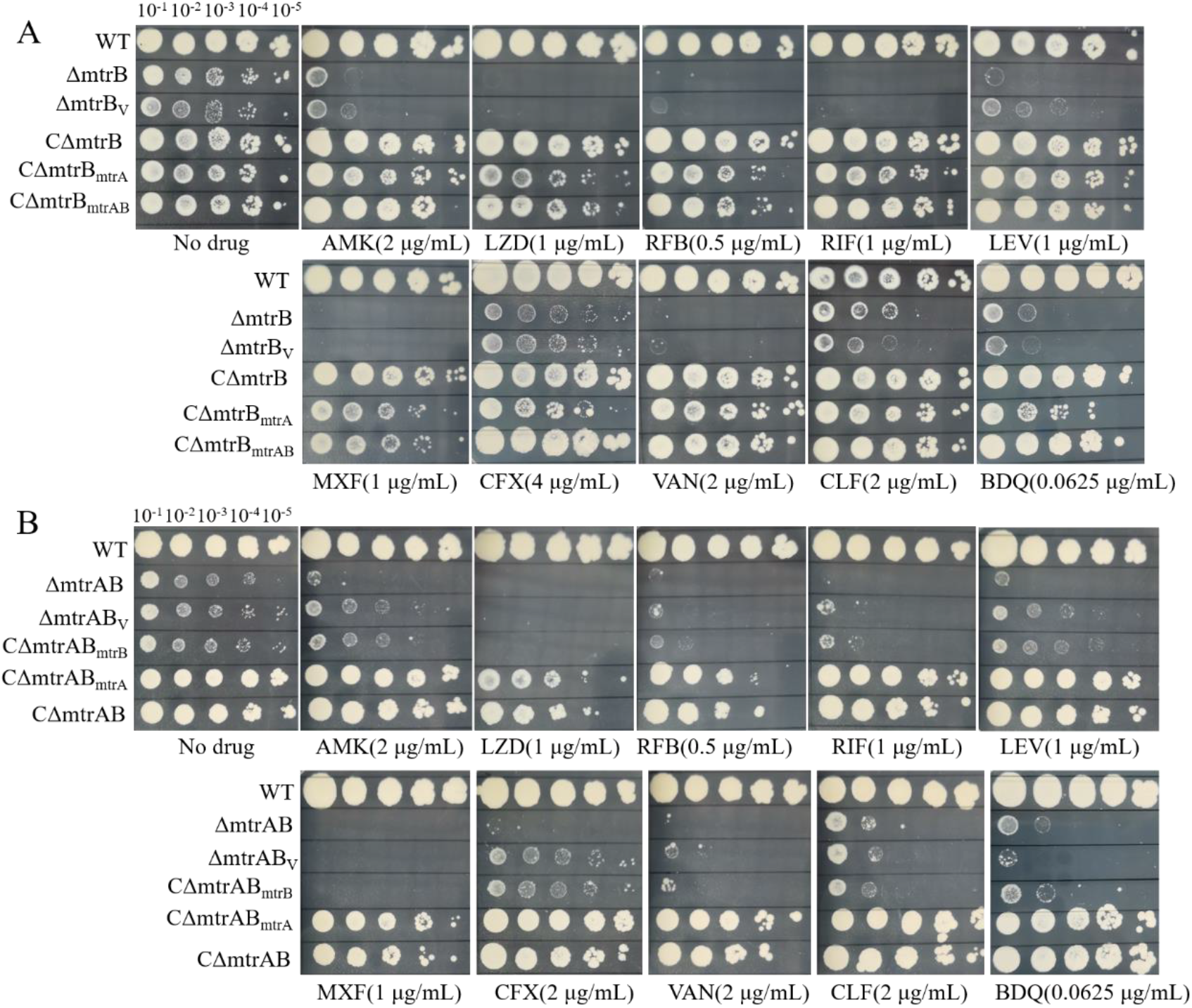
Deletion of *mtrB* (*MAB_3590c*) or *mtrB* plus *mtrA* (*MAB_3590c-MAB_3591c*) confers heightened sensitivity to various drugs. (A)The susceptibility measurements for WT (the wild-type Mab), ΔmtrB (*mtrB* knockout Mab), ΔmtrB_V_ (ΔmtrB carrying pMV261 empty plasmid), CΔmtrB (ΔmtrB carrying pMV261-mtrB expressing *mtrB* under the strong mycobacterial promoter *hsp60*), CΔmtrB_mtrA_ (ΔmtrB carrying pMV261-mtrA expressing *mtrA*), CΔmtrB_mtrAB_ (ΔmtrB carrying pMV261-mtrAB expressing *mtrAB*) against different drugs in plates. (B) The susceptibility measurements for WT (the wild-type Mab), ΔmtrAB (*mtrAB* knockout strain), ΔmtrAB_V_ (ΔmtrAB carrying pMV261 empty plasmid), CΔmtrAB_mtrB_ (ΔmtrAB carrying pMV261-mtrB expressing *mtrB*), CΔmtrAB_mtrA_ (ΔmtrAB carrying pMV261-mtrA expressing *mtrA*), and CΔmtrAB (ΔmtrAB carrying pMV261-mtrAB expressing *mtrAB*) against different drugs in plates.

**Table 2.** MICs of various drugs against different Mab strains.

| Drugs | Mab strains/ MICs ($\mu\text{g/mL}$) | | | | | |
| --- | --- | --- | --- | --- | --- | --- |
| | WT | $\Delta mtrB$ | $\Delta mtrB_V$ | $C\Delta mtrB$ | $C\Delta mtrB_{mtrA}$ | $C\Delta mtrB_{mtrAB}$ |
| Amikacin | 16 | 1 | 1 | 4 | 4 | 4 |
| Linezolid | 64 | 0.5-1 | 0.5-1 | 4 | 4 | 4 |
| Rifabutin | 4 | 0.25 | 0.25 | 1 | 1 | 1 |
| Rifampin | >128 | 2 | 1-2 | 16 | 8 | 8 |
| Levofloxacin | 32 | 2-4 | 2 | 8 | 4 | 2 |
| Moxifloxacin | 8-16 | 2-4 | 2 | 8 | 8 | 8 |
| Cefoxitin | 32 | 2-4 | 4 | 16 | 16 | 16 |
| Vancomycin | 32 | 1 | 1 | 8 | 4 | 4 |
| Clofazimine | 4 | 1 | 0.5 | 2 | 2 | 2 |
| Bedaquiline | 0.5 | 0.0625 | 0.0625 | 0.25 | 0.25 | 0.25 |

WT, the wild-type Mab; $\Delta mtrB$ , *mtrB* knockout Mab; $\Delta mtrB_V$ , $\Delta mtrB$ carrying pMV261 empty plasmid; $C\Delta mtrB$ , $\Delta mtrB$ carrying pMV261-*mtrB* expressing *mtrB* under the strong mycobacterial promoter *hsp60*; $C\Delta mtrB_{mtrA}$ , $\Delta mtrB$ carrying pMV261-*mtrA* expressing *mtrA*; $C\Delta mtrB_{mtrAB}$ , $\Delta mtrB$ carrying pMV261-*mtrAB* expressing *mtrAB*.

**Table 3.** MICs of various drugs against different Mab strains.

| Drugs | Mab strains/ MICs ($\mu\text{g/mL}$) | | | | | |
| --- | --- | --- | --- | --- | --- | --- |
| | WT | $\Delta mtrAB$ | $\Delta mtrAB_V$ | $C\Delta mtrAB_{mtrB}$ | $C\Delta mtrAB_{mtrA}$ | $C\Delta mtrAB$ |
| Amikacin | 16 | 1 | 4 | 4 | 16 | 16 |
| Linezolid | 64 | 0.5-1 | 1 | 1 | 8 | 16 |
| Rifabutin | 4 | 0.125 | 0.125 | 0.25 | 4 | 4 |
| Rifampin | >128 | 0.25 | 0.25 | 0.5 | 32 | 64 |
| Levofloxacin | 32 | 0.5 | 0.5 | 1 | 8 | 4 |
| Moxifloxacin | 8-16 | 0.25 | 0.25 | 0.5 | 8 | 4 |
| Cefoxitin | 32 | 1 | 1 | 1 | 4 | 8 |
| Vancomycin | 32 | 0.25 | 0.25 | 0.5 | 8 | 4 |
| Clofazimine | 4 | 0.5 | 0.5 | 1 | 2 | 2 |
| Bedaquiline | 0.5 | 0.0625 | 0.0625 | 0.25 | 0.0625 | 0.0625 |

WT, the wild-type Mab; $\Delta mtrAB$ , *mtrAB* knockout Mab; $\Delta mtrAB_V$ , $\Delta mtrAB$ carrying pMV261 empty plasmid; $C\Delta mtrAB_{mtrB}$ , $\Delta mtrAB$ carrying pMV261-*mtrB* expressing *mtrB*; $C\Delta mtrAB_{mtrA}$ , $\Delta mtrAB$ carrying pMV261-*mtrA* expressing *mtrA*; $C\Delta mtrAB$ , $\Delta mtrAB$ carrying pMV261-*mtrAB* expressing *mtrAB*.

### 3.4 Deletion of *mtrA* or *mtrB* sensitizes Mab to multiple drugs *in vivo*

Given the elevated drug susceptibility of knockout strains *in vitro*, we questioned whether they could become susceptible to previously ineffective drugs against wild-type Mab *in vivo*. We next examined the activities of drugs including AMK, BDQ, RFB, MXF and LZD against knockout strains in mice infected with WT, ΔmtrA, and ΔmtrB. After a 10-day treatment period, in ΔmtrA infected mice, AMK, BDQ, and RFB significantly reduced bacterial burden in the lungs compared to the vehicle group (*p*< 0.01) (Figure. 3). In the ΔmtrB infected mice, AMK and BDQ showed similar activities, while RFB led to an even more significant bacterial decrease (*p*<0.001). However, the other two drugs, MXF and LZD, did not exhibit noticeable activities (Figure. 3B). Hence, AMK, BDQ and RFB were active against ΔmtrA and ΔmtrB *in vivo* at the tested doses, suggesting the potential of MtrA and MtrB as drug targets.

**Figure 3.**
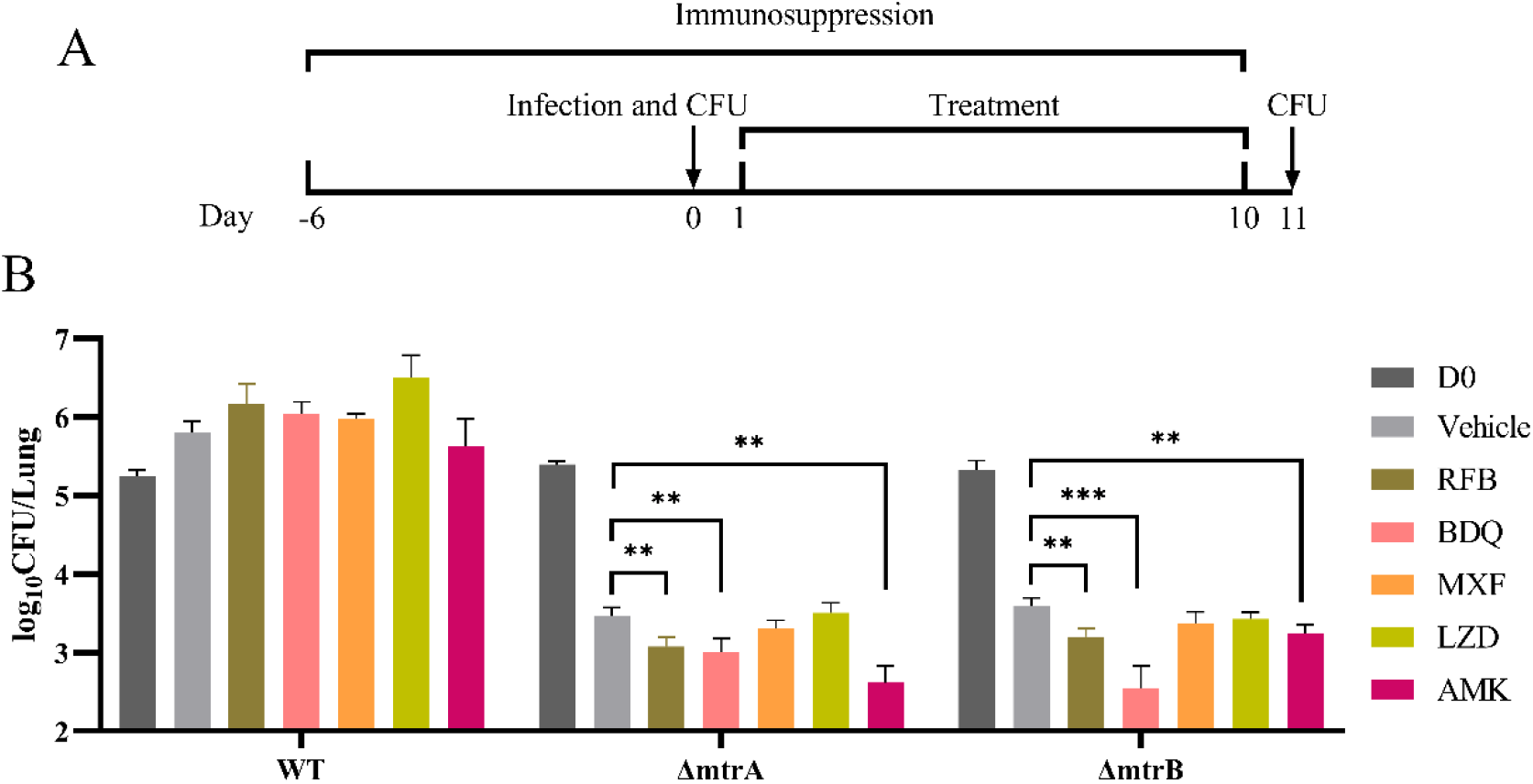
Assessment of susceptibility to antibiotics *in vivo*. (A) Schematic representation of the murine Mab lung infection model used in this study. All mice were immunosuppressed with DEXA 7 days prior to infection and remained so throughout the experiment. Lungs were homogenized and plated on agar for CFU determination at three hours after infection and on one day following the completion of treatment. Treatment commenced on the first day post-infection and continued for ten days. (B) Comparison of Mab CFUs in the lungs of mice treated with antibiotic versus untreated controls. Drugs were administered at the following doses (mg/kg): RFB 10, BDQ 20, MXF 100, LZD 100, AMK 150. Unpaired *t*-test were used for statistical significance. **, *p* < 0.01; ***, *p* < 0.001.,

### 3.5 Deletion of *mtrA, mtrB* or both significantly diminishes the virulence of Mab *in vivo*

The bacterial load in mice infected with the knockout strains, without treatment, significantly decreased compared to the load at the initiation of treatment. This reduction is likely attributed the attenuated virulence of the bacteria. Six days post-infection, the number of WT bacteria recovered from the lungs increased from 5.23 ± 0.10 log_10_ to 5.97 ± 0.12 log_10_ (mean ± standard deviation [SD]). Conversely, the bacterial burden of the three knockout strains exhibited varying degrees of reduction (Figure. 4 and Table. 4). The CFUs of ΔmtrA and ΔmtrB in the lungs demonstrated a consistent decrease, from 5.40 ± 0.04 log_10_ to 4.34 ± 0.02 log_10_, and from 5.31 ± 0.15 log_10_ to 4.35 ± 0.10 log_10_, respectively. However, the double knockout strain ΔmtrAB showed a marked decrease, from 4.71 ± 0.06 log_10_ to 2.98 ± 0.11 log_10_. By day 11, the count of WT had stabilized while the counts of ΔmtrA and ΔmtrB had dropped to 3.45 ± 0.11 log_10_ and 3.59 ± 0.11 log_10_ respectively. Notably, ΔmtrAB showed a pronounced reduction to 2.06 ± 0.25 log_10_ (less than 100 CFU/lung). By day 16, the bacterial load of knockout strains in the lungs continued to decline, with ΔmtrAB showing a near 3 log_10_ reduction, reaching 1.69 ± 0.40 log_10_. In summary, these results indicate that MtrAB plays a critical role in the virulence of Mab and highlight the requirement of MtrAB for sustaining infection in an *in vivo* model. The drastic attenuation of virulence further highlights the promise of MtrA and MtrB as pharmaceutical targets.

**Figure 4.**
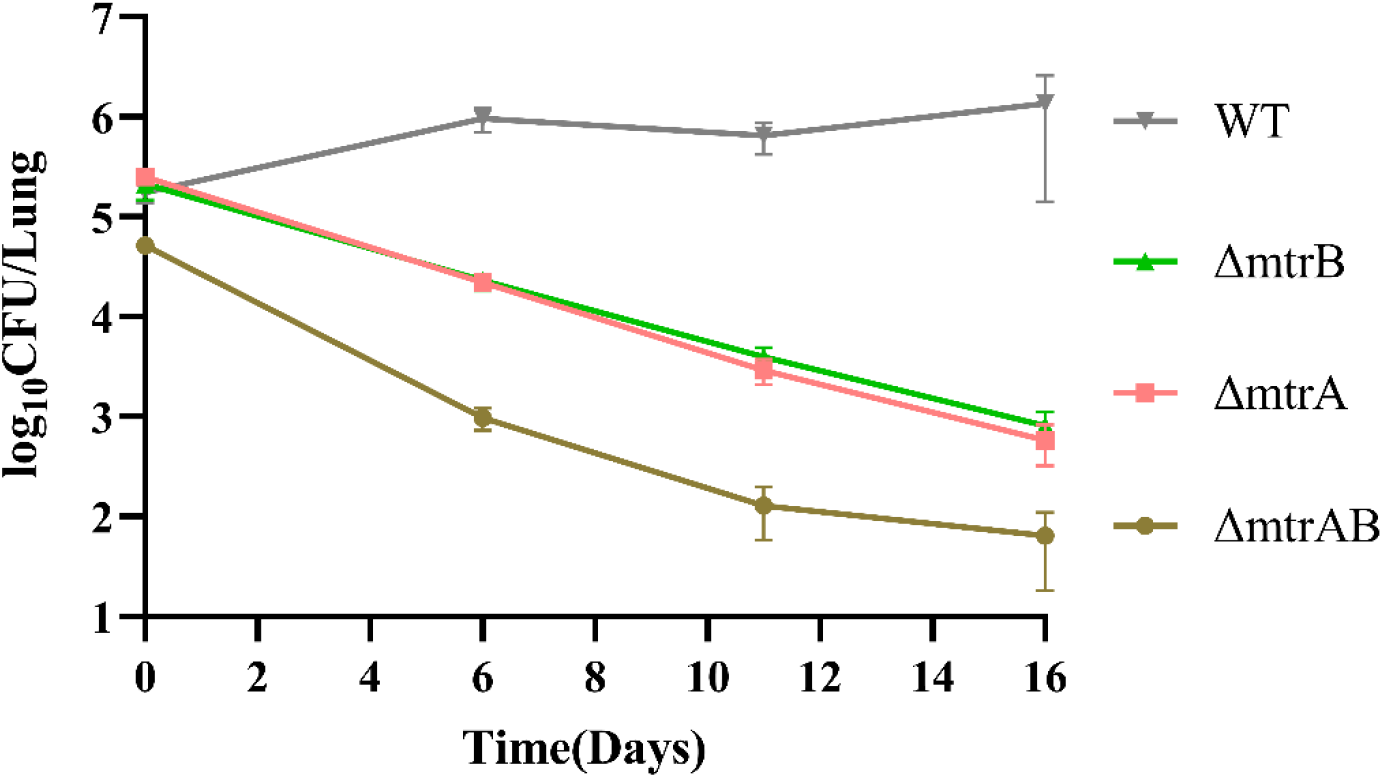
Growth curves of different Mab strains in a mouse model. WT, the wild-type Mab; ΔmtrA, *mtrA* knockout Mab; ΔmtrB, *mtrB* knockout Mab; ΔmtrAB, *mtrAB* knockout Mab. D0 denotes the day on which the initial infection bacterial load is examined.

**Table 4.** Bacterial burden in the lungs of mice infected with various Mab strains in the indicated time points.

| Group | Mean ± SD log<sub>10</sub> CFU/Lung | | | |
| --- | --- | --- | --- | --- |
|  | D0 | D6 | D11 | D16 |
| WT | 5.23 ± 0.10 | 5.97 ± 0.12 | 5.79 ± 0.17 | 5.95 ± 0.49 |
| $\Delta mtrA$ | 5.40 ± 0.04 | 4.34 ± 0.02 | 3.45 ± 0.11 | 2.74 ± 0.18 |
| $\Delta mtrB$ | 5.31 ± 0.15 | 4.35 ± 0.10 | 3.59 ± 0.11 | 2.89 ± 0.14 |
| $\Delta mtrAB$ | 4.71 ± 0.06 | 2.98 ± 0.11 | 2.06 ± 0.25 | 1.69 ± 0.40 |

Results are presented as mean ± standard deviation (SD) log<sub>10</sub> CFU/Lung. D0 denotes the day of assessment for the initial infection bacterial load. WT, the wild-type Mab; $\Delta mtrA$ , *mtrA* knockout Mab; $\Delta mtrB$ , *mtrB* knockout Mab; $\Delta mtrAB$ , *mtrAB*

### 3.6 Deletion of either *mtrA, mtrB* or both impacts the growth and cell morphology of Mab

In investigating the factors contributing to reduced virulence of the knockout strains and their enhanced response to durgs *in vivo*, we noted that MtrA in Mtb was implicated in cell division and cell wall metabolism[30, 31]. We observed that ΔmtrA, ΔmtrB and ΔmtrAB all exhibited significantly reduced growth rates compared to WT (Figure. 5). Gene reintroduction into these strains partially restored growth, indicating that the MtrAB is crucial for maintaining growth.

**Figure 5.**
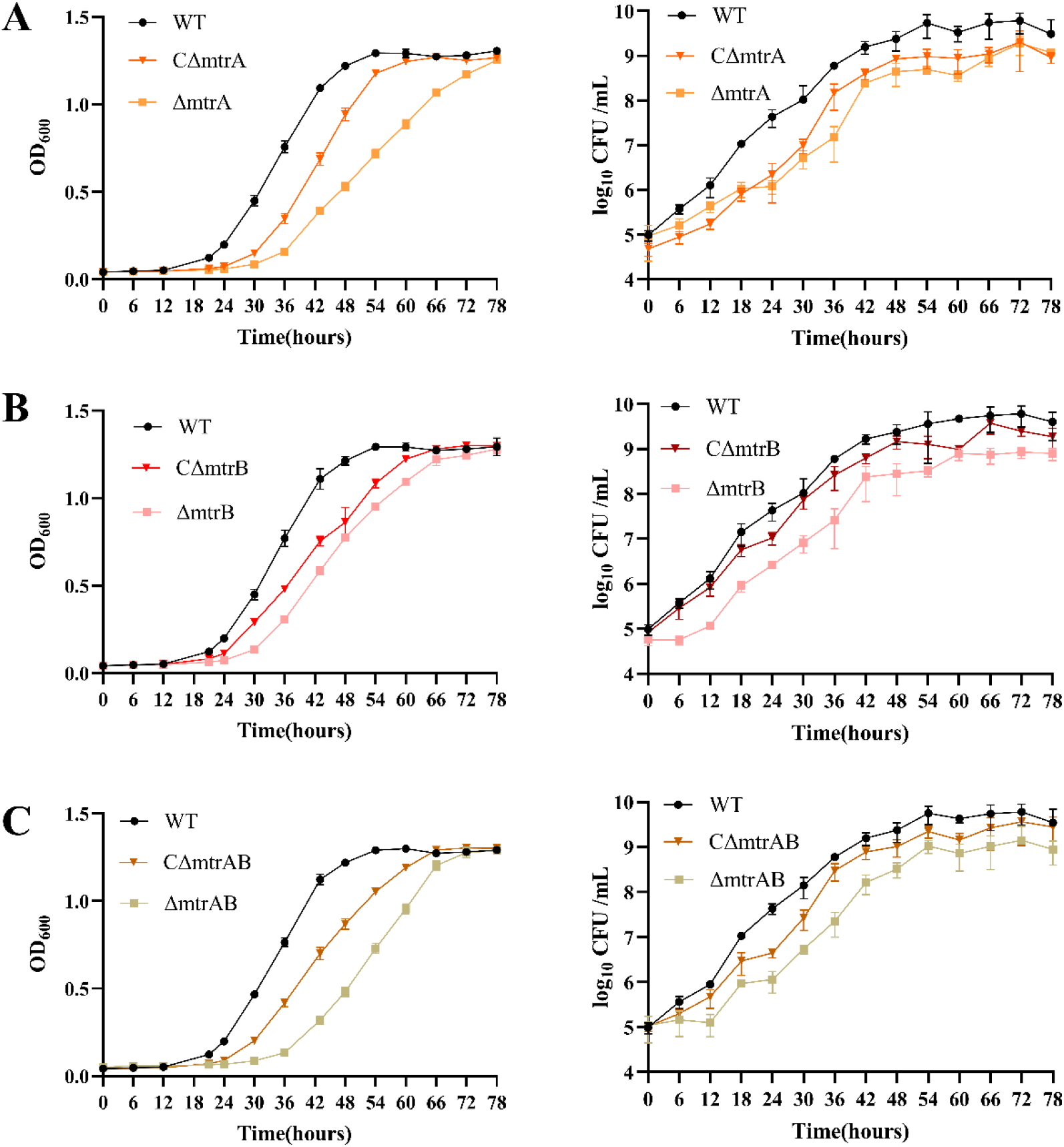
The deletion of either *mtrA, mtrB* or both results in a slower growth rate for Mab. The growth curves (shown on the left by OD_600_ and shown on the right by CFUs) are depicted for the *mtrA* (A), *mtrB* (B), and *mtrAB* (C) knockout strains, as well as their respective complementary strains. WT, the wild-type Mab; ΔmtrA, *mtrA* knockout Mab; CΔmtrA, ΔmtrA carrying pMV261-mtrA expressing *mtrA*; ΔmtrB, *mtrB* knockout Mab; CΔmtrB, ΔmtrB carrying pMV261-mtrB expressing *mtrB*; ΔmtrAB, *mtrAB* knockout Mab; CΔmtrAB, ΔmtrAB carrying pMV261-mtrAB expressing *mtrAB*.

Subsequently, we examined the morphology of ΔmtrA, ΔmtrB, ΔmtrAB and WT as well as the respective complementary strains CΔmtrA, CΔmtrB, CΔmtrAB, Significant morphological changes were evident in the knockout strains (Figure. 6A). The knockout strains were notably elongated with average lengths of ΔmtrA (3.6 ± 1.11 mm), ΔmtrB (3.59 ± 0.85 mm), and ΔmtrAB (4.53 ± 1.03 mm), exceeding that of the wild-type (2.18 ± 0.37 mm) (Figure. 6B). Gene complementation restored cell lengths comparable to WT. In addition, these strains formed multiseptated chains more frequently than WT and complementary strains (Figure. 6C), indicating a defect in cell division. It appears that Mab lacking MtrAB undergoes normal elongation and septum formation, but falters in the separation into daughter cells. These findings indicate that MtrAB is involved in Mab cell division.

**Figure 6.**
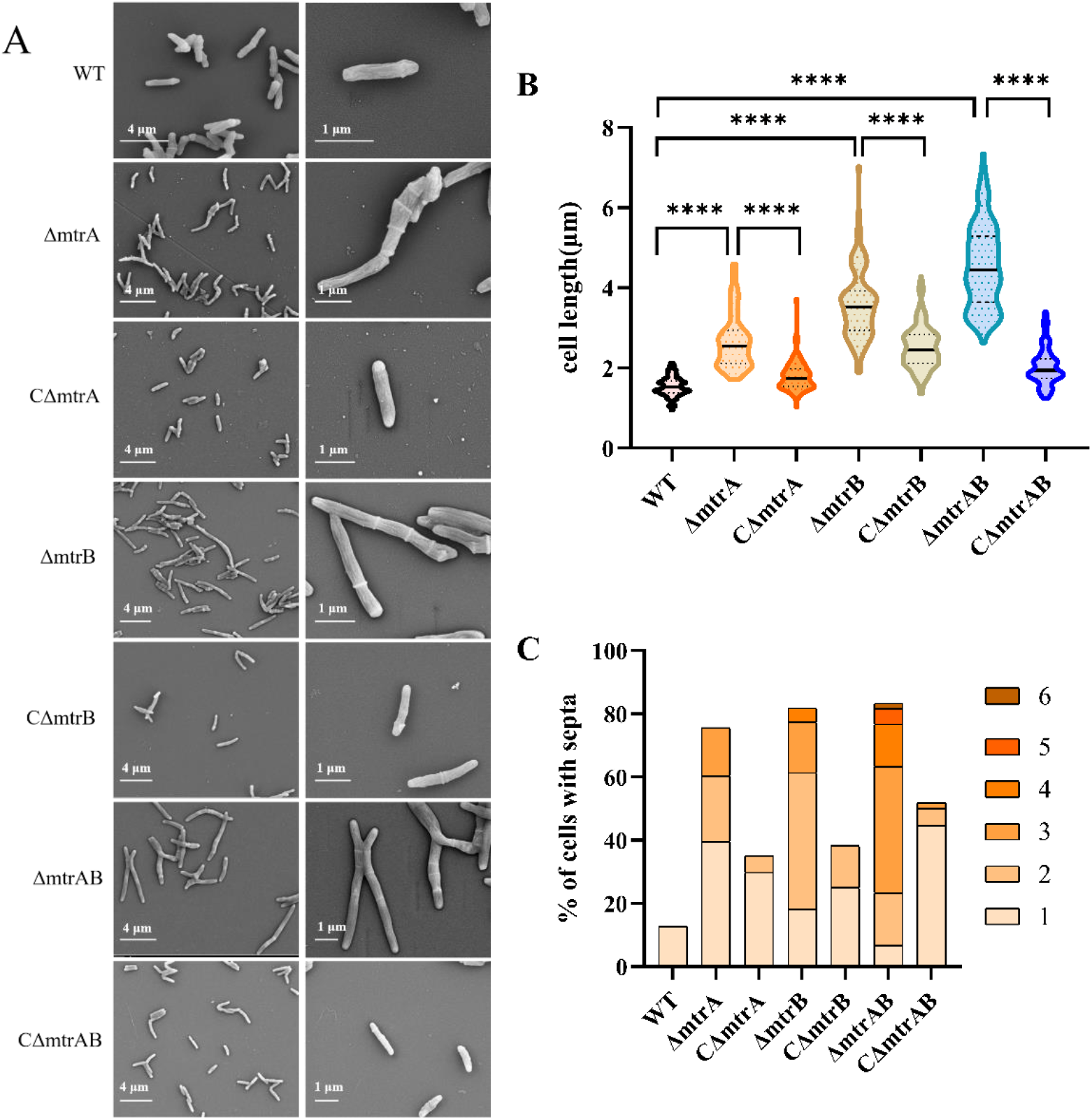
The impact of deleting *mtrA, mtrB*, or both on Mab cell morphology. (A) Representative micrographs of the knockout and complementary strains captured via cryo-scanning electron microscopy. (B) Violin plots depict the distribution of cell lengths for the wild-type Mab (WT; n = 111), knockout Mab strains (ΔmtrA; n = 114, ΔmtrB; n = 98, and ΔmtrAB; n = 109), and the corresponding complementary strains (CΔmtrA; n = 108, CΔmtrB; n = 109, and CΔmtrAB; n = 109). The median is denoted by a black line, with the interquartile range indicated by dashed lines. Statistical significance was determined using an unpaired *t*-test, with asterisks (****, *p* < 0.0001) indicating significance. (C) Quantitative analysis of the percentage of cells containing multiple septa in the wild-type (WT; n = 101) and various knockout (ΔmtrA; n = 106, ΔmtrB; n = 88, and ΔmtrAB; n = 120) and complementary strains (CΔmtrA; n = 114, CΔmtrB; n = 120, and CΔmtrAB; n = 112). WT, the wild-type Mab; ΔmtrA, *mtrA* knockout Mab; CΔmtrA, ΔmtrA carrying pMV261-mtrA expressing *mtrA*; ΔmtrB, *mtrB* knockout Mab; CΔmtrB, ΔmtrB carrying pMV261-mtrB expressing *mtrB*; ΔmtrAB, *mtrAB* knockout Mab; CΔmtrAB, ΔmtrAB carrying pMV261-mtrAB expressing *mtrAB*.

### 3.7 Deletion of either *mtrA, mtrB* or both resulted in an increased cell envelope permeability of Mab

Cell elongation during the initial phase of cell division, along with the formation and subsequent separation of the septum in the later stages, is characterized by a reorganization of cell envelope constituents[32]. To assess envelope integrity, we measured permeability using an EtBr uptake assay. Notably, Mab strains deficient in either MtrA, MtrB, or both MtrA and MtrB exhibited accelerated and augmented accumulation of EtBr compared to WT (Figure. 7). The complementation mitigated the accumulation partially but significantly reduced the EtBr accumulation. Collectively, these findings point to increased permeability of the knockout strains’ cell envelopes, and indirectly confirm to the MtrAB’s regulatory role in the cell division process. Interestingly, we observed that the knockout strains exhibited higher sedimentation rates than WT (Figure. S3), they were more prone to forming bacterial clumps, which may be attributed to alterations in cell wall composition affecting bacterial aggregation.

**Figure 7.**
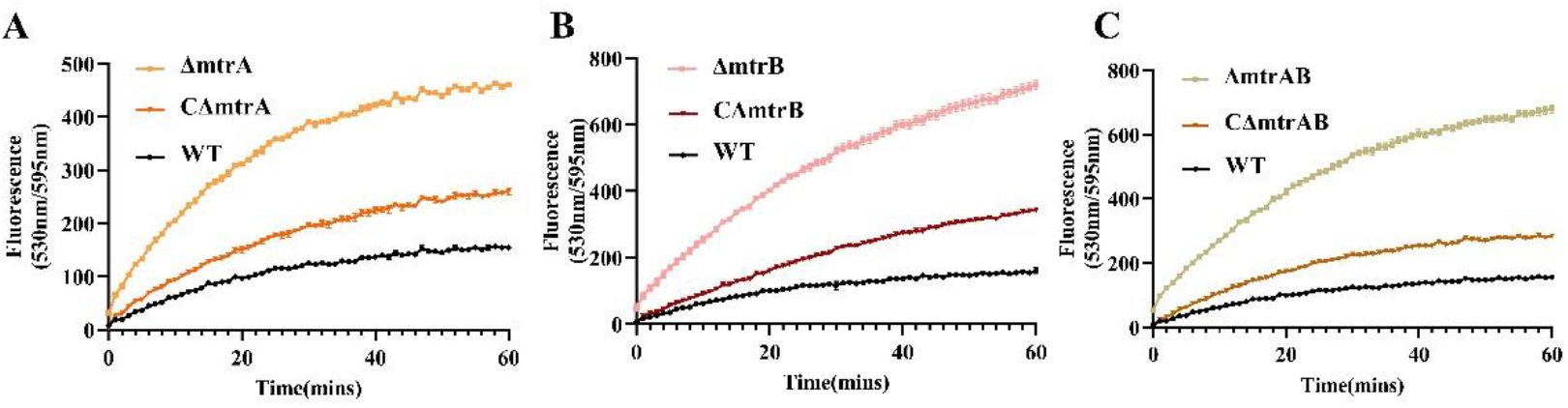
The deletion of *mtrA, mtrB* or both in Mab leads to increased cell envelope permeability. The fluorescence, which signifies the accumulation of EtBr in various strains, was monitored for the *mtrA* (A), *mtrB* (B), and *mtrAB* (C) knockout strains, as well as their respective complementary strains, over a period of 60 minutes. WT, the wild-type Mab; ΔmtrA, *mtrA* knockout Mab; CΔmtrA, ΔmtrA carrying pMV261-mtrA expressing *mtrA*; ΔmtrB, *mtrB* knockout Mab; CΔmtrB, ΔmtrB carrying pMV261-mtrB expressing *mtrB*; ΔmtrAB, *mtrAB* knockout Mab; CΔmtrAB, ΔmtrAB carrying pMV261-mtrAB expressing *mtrAB*.

## 4. Discussion

Mab have recently emerged as an important opportunistic pathogen, causing serious lung diseases and skin and soft tissue infections[1]. It is known as an “incurable nightmare” due to the considerable challenges it presents in treatment[5, 6]. The underlying cause of this challenge is its intrinsic drug resistance to a wide array of drugs. In this study, we discovered that the MtrAB TCS encoded by *MAB_3591c(mtrA)* and *MAB_3590c(mtrB)*, influences both the intrinsic resistance and virulence of Mab through the regulation of cell division. Consequently, MtrA and MtrB represent potential targets for the development of anti-Mab drugs, which could not only enhance the efficacy of other drugs but also augment the killing efficacy of the host immune system.

In this study, we have preliminarily demonstrated that the disruption of either *mtrA, mtrB*, or both in Mab, significantly impacts intrinsic resistance to drugs with diverse mechanisms of action (Figure. 1 and Figure. 2). This includes translation inhibitors, cell envelope synthesis inhibitors, fluoroquinolones that disrupt DNA replication, as well as respiratory chain targeting drugs. The increased drug susceptibility may result from the compromised cell envelope, which allows for the enhanced penetration of various antimicrobial agents through what was previously a robust barrier, thus increasing their effectiveness. This hypothesis is supported by the discovery of increased permeability and altered cell morphology in the knockout strains (Figure. 6 and Figure. 7). Conversely, in Mtb, *mtrA* knockdown renders Mtb more susceptible to drugs tested with molecular weights above 200 g/mol; however, but there is no change in sensitivity to the translation inhibitor LZD[18, 33]. This indicates a distinctive influence of the MtrAB TCS on drug resistance mechanisms between these mycobacterial species.

The virulence traits of Mab render it pathogenic in fragile hosts with structural lung disease. The interactions between the robust mycobacterial lipid-rich sturdy cell envelope and host immune cells enables Mab to survive within the host’s challenging internal environment[34–36]. Several identified virulence factors (VFs) are associated with cell envelope. Mycobacterial membrane protein large (MmpL) proteins are a family of VFs of the Mycobacterium genus, crucial for transporting lipids vital to maintaining the integrity of the mycobacterial envelope and for delivering siderophores to the periplasmic space[37]. Additionally, research has indicated that the clustering of Mab glycopeptidolipids on specific nanodomains of the bacterial surface regulates surface hydrophobicity, thereby influencing bacterial adhesion and virulence[38]. In this study, we found that strains with knockouts of *mtrA, mtrB*, or both displayed reduced virulence *in vivo* (Figure. 4 and Table. 4). From this observation, we speculate that MtrAB may influence cell virulence by regulation of cell envelope rearrangement during cell division. Nevertheless, the direct regulation of a particular virulence factor by MtrAB cannot be dismissed and warrants further investigation. The pronounced decrease in virulence exhibited by the knockout strains implies that MtrA and MtrB are attractive candidate targets for drug development.

The deletion of either *mtrA* or *mtrB* resulted in heightened susceptibility to RFB, AMK, and BDQ *in vivo* (Figure. 3B), reinforcing the concept that MtrA and MtrB could serve as viable drug targets. Conversely, MXF and LZD exhibited no *in vivo* activity against the knock strains, likely due to their bacteriostatic rather than bactericidal properties in Mab[39, 40]. Moreover, a slight rise in bacterial load (from 5.79 ± 0.17 log_10_ to 6.23 ± 0.67 log_10_) was noted in mice infected with WT and treated with LZD, which may be attributed to the toxic impact of LZD on the host, possibly promoting bacterial growth. Nevertheless, mice infected with the knockout strains and treated with LZD did not exhibit an increase in bacterial load compared to the untreated control, suggesting the potential bacteriostatic effectiveness of LZD.

We noted that the overexpression of *mtrA* in both ΔmtrB and ΔmtrAB successfully restored drug resistance. The strains CΔmtrB_mtrA_ and CΔmtrAB_mtrA_, which lack *mtrB* and overexpress *mtrA*, exhibited a restored phenotype, contrasting with the elevated drug susceptibility observed in ΔmtrB. Several hypotheses could explain this outcome. One possibility is that the expression of *mtrA* is controlled by MtrB, without *mtrB, mtrA* is not expressed in the ΔmtrB strain. However, in both CΔmtrB_mtrA_ and CΔmtrAB_mtrA_ strains, where *mtrA* is constitutively expressed under the *hsp60* promoter, it can function effectively. Another possibility is that MtrA may be phosphorylated by alternative kinases or undergo autophosphorylation, yet this requires a specific MtrA threshold to trigger the reaction, which may not have been met in the ΔmtrB strain. Furthermore, there might be other mechanisms regulating *mtrA* that are independent of MtrB. These assumptions require additional scrutiny.

Upon creating the double knockout strain ΔmtrAB, we noticed that it had increased drug sensitivity and decreased virulence compared to either ΔmtrA or ΔmtrB. We hypothesize that this heightened sensitivity and reduced virulence upon deletion of the entire TCS may stem from the modular nature of two-component protein domains, allowing for varied integration into different proteins and pathways[41]. Without MtrA, MtrB could potentially phosphorylate alternate effector (s), preserving its role in drug resistance. Similarly, without MtrB, MtrA might be phosphorylated by other kinases, enabling it to continue functioning. Yet, when the entire TCS is deleted, the expression of many genes related to drug resistance is hindered, resulting in heightened sensitivity. This is corroborated by evidence showing that MtrA acts as a substrate of protein kinase PknK in Mtb, which can phosphorylate it, highlighting the intricate interaction within the TCS[42].

## 5. Conclusions

Our study underscores the pivotal role of MtrAB TCS in conferring intrinsic resistance and virulence of Mab through its regulation of cell division. These findings compellingly indicate that MtrAB could be an ideal target for the development of novel anti-Mab drugs.

## Supporting information

Graphical Abstract

