## Supplementary for "MtrAB two-component system is crucial for the intrinsic resistance and virulence of *Mycobacterium abscessus*"

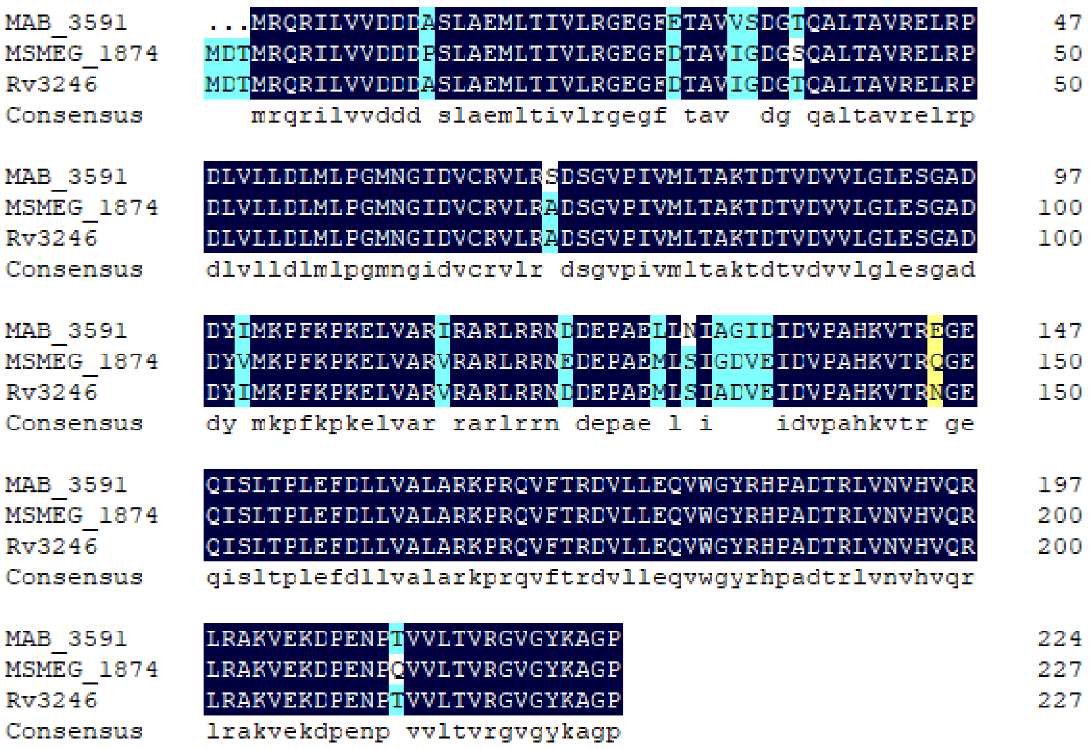


**Figure S1.** The amino acid sequences of MtrA in Mab (MAB_3591), Mtb (Rv3246), and Msm (MSMEG_1874) were aligned. The MtrA sequence exhibited a 93.86% identity with that of Mtb and a 91.23% identity with the sequence from Msm.

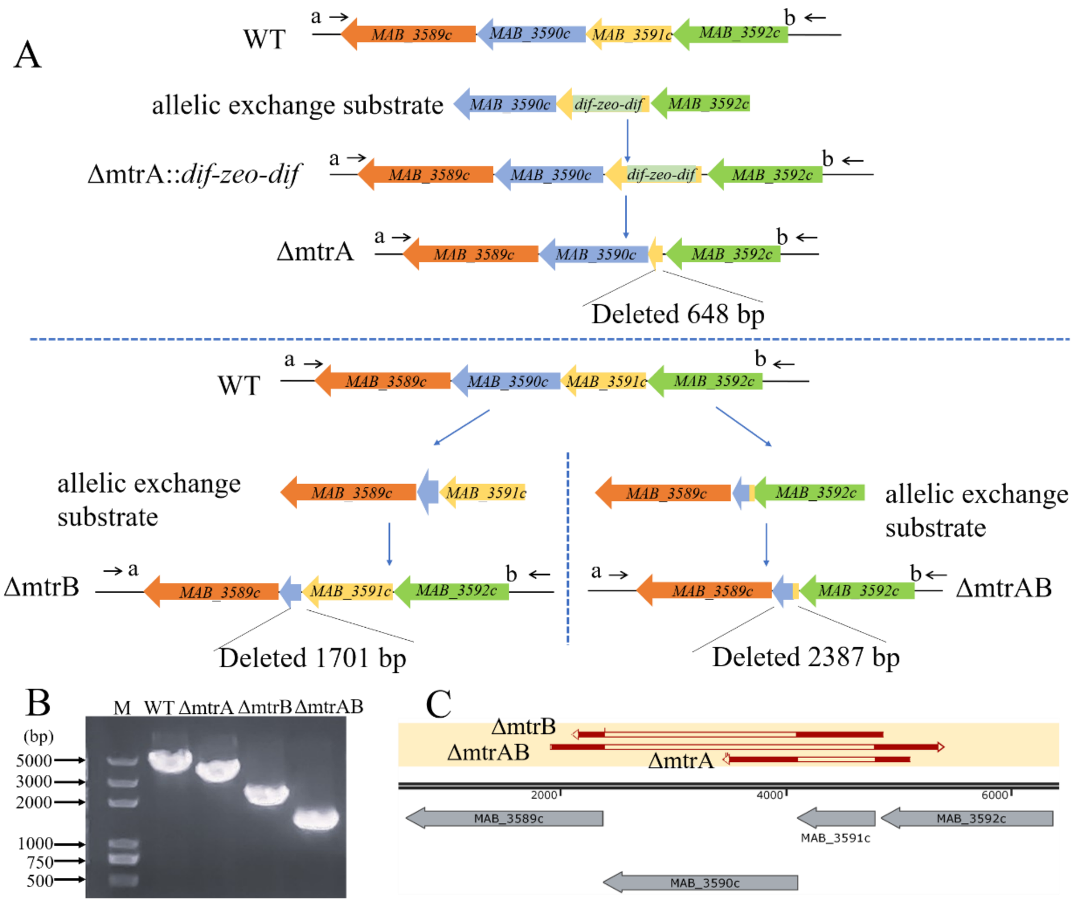


**Figure S2.** (A) Schematic representation of the construction of selectable marker-free knockout strains ΔmtrA(upper), ΔmtrB (bottom left), ΔmtrAB (bottom right). (B)PCR identification results for the knockout strains are shown. The primers used for identification are labeled as a and b in (A). Lane M contains the DNA marker, while the remaining lanes, from left to right, display the PCR products for the wild type (WT), *mtrA* knockout strain ΔmtrA, *mtrB* knockout strain ΔmtrB, and *mtrAB* knockout strain ΔmtrAB. The observed patterns align with the predicted fragment sizes: wild type at 3.7 kb, knockout fragments at 3.1 kb, 2.0 kb, and 1.1 kb. (C) Sequence alignment results for the *MAB_3591c-MAB_3590c* genes among the knockout strains and the wild-type strain are illustrated. The gap in the alignment indicates the deleted gene fragment.

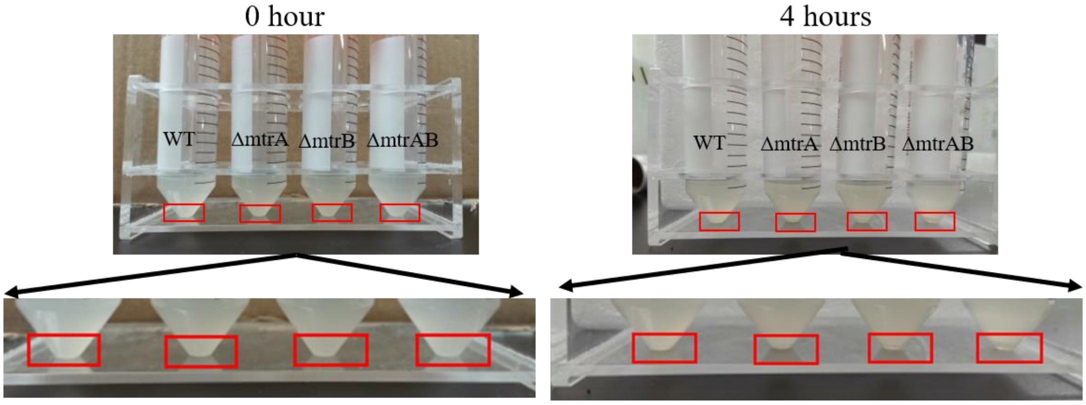


**Figure S3.** Comparison of wild-type Mab and *mtrA*, *mtrB*, *mtrAB* knockout Mab strains bacterial sedimentation rates. Photographs were captured following a static settling period of 0 and 4 hours for the bacterial culture, which remained in an identical condition. WT, wild-type Mab; ΔmtrA, *mtrA* knockout Mab; ΔmtrB, *mtrB* knockout Mab; ΔmtrAB, *mtrAB* knockout Mab.

**Table S1.** Strains used in this study

| **Strains** | **Source or reference** |
|---|---|
| *Mycobacterium abscessus* GZ002 | [1] |
| I5 | [2] |
| ΔmtrA | This study |
| CΔmtrA | This study |
| ΔmtrB | This study |
| CΔmtrB | This study |
| CΔmtr$B_{mtrA}$ | This study |
| CΔmtr$B_{mtrAB}$ | This study |
| ΔmtrAB | This study |
| CΔmtrAB | This study |
| CΔmtr$AB_{mtrA}$ | This study |
| CΔmtr$AB_{mtrB}$ | This study |

**Table S2.** Plasmids used in this study

| **Plasmids** | **Source or reference** |
|---|---|
| pMV261 | [3] |
| pMV261-mtrA | This study |
| pMV261-mtr$A_{Ms}$ | This study |
| pMV261-mtr$A_{Mtb}$ | This study |
| pMV261-mtrB | This study |
| pMV261-mtrAB | This study |
| pCR-Zeo | [4] |
| pCR-Zeo-mtrB | This study |

| | |
|---|---|
| pBluescript II SK(+) | [5] |
| pBluescript II SK(+)-mtrA-UZD | This study |
| pBluescript II SK(+)-mtrB-UD | This study |
| pBluescript II SK(+)-mtrAB-UD | This study |
| pJV53 | [6] |
| pJV53-Cpf1 | [4] |

**Table S3.** Oligonucleotides used in this study

| **Name** | **Sequence (5’-3’)** | **Description** |
|---|---|---|
| mtrB-crRNA-F | ATGTGGCTCACATCAGAGGTGAAGA | crRNA for *mtrB* and *mtrAB* knockout |
| mtrB-crRNA-R | AGCTTCTTCACCTCTGATGTGAGCCACATCT | |
| cz-mtrBUP-F | TCCGCAAGCTtGCTGAAGATCACGGGGGTCC | Construct a plasmid containing *mtrB* upstream and downstream homologous arms |
| cz-mtrBUP-R | CGCTCTAGAACTAGTGGATCCTGCTCACCATCGTGTTGCG | |
| cz-mtrBDOWN-F | CTCGAGGTCGACGGTATCGATAGTTCGGTGGCCAGCAGAT | |
| cz-mtrBDOWN-R | ATCTTCAGCaAGCTTGCGGAGGATGCG | |
| mtrB-UD-F | AGTTCGGTGGCCAGCAGAT | Amplify upstream and downstream homologous arms fragment of *mtrB* |
| mtrB-UD-R | TGCTCACCATCGTGTTGCG | |
| cz-mtrABUP-F | CAGATCCCATCGTGTCACTAGATCG | Construct a plasmid containing *mtrAB* upstream and downstream homologous arms |
| cz-mtrABUP-R | CGCTCTAGAACTAGTGGATCCGATGTGCAGGAGCAGGCCC | |
| cz-mtrABDOWN-F | CTCGAGGTCGACGGTATCGATAGTTCGGTGGCCAGCAGAT | |
| cz-mtrABDOWN-R | TAGTGACACGATGGGATCTGCGGAGGATGCGTCGTGA | |
| mtrAB-UD-F | AGTTCGGTGGCCAGCAGAT | Amplify upstream and downstream homologous arms fragment of *mtrAB* |
| mtrAB-UD-R | GATGTGCAGGAGCAGGCCC | |
| JD-KOAB-F | GGGGAGAATGTCACCTCCA | identify the knockout of *mtrA*, *mtrB* and *mtrAB* |
| JD-KOAB-R | CCCGTTCAACAATCCGTAG | |
| cz-mtrB-F | GGCCAAGACAATTGCGGATCCGTGATCTTCAGCTCGAAGCGG | Construct a plasmid expressing *mtrB* from Mab |
| cz-mtrB-R | GTTAACTACGTCGACATCGATTCACGACGCATCCTCCGC | |
| cz-mtrAB-F | GGCCAAGACAATTGCGGATCCATGGGATCTATGAGGCAAAGGA | Construct a plasmid expressing *mtrAB* from Mab |
| cz-mtrAB-R | GTTAACTACGTCGACATCGATTCACGACGCATCCTCCGC | |
| cz-mtrA-Ms-F | GGCCAAGACAATTGCGGATCCATGGACACCATGAGGCAAAGG | Construct a plasmid expressing *mtrA* from Msm |
| cz-mtrA-Ms-R | GTTAACTACGTCGACATCGATTCACGGGGGTCCGGCCTT | |
| cz-mtrA-Mtb-F | GGCCAAGACAATTGCGGATCCATGAGGCAAAGGATTTTGGTCG | Construct a plasmid expressing *mtrA* from Mtb |
| cz-mtrA-Mtb-R | GTTAACTACGTCGACATCGATTCACGGAGGTCCGGCCTT | |
