## Supplementary figures and images for "MtrAB two-component system is crucial for the intrinsic resistance and virulence of *Mycobacterium abscessus*"

### Graphical Abstract

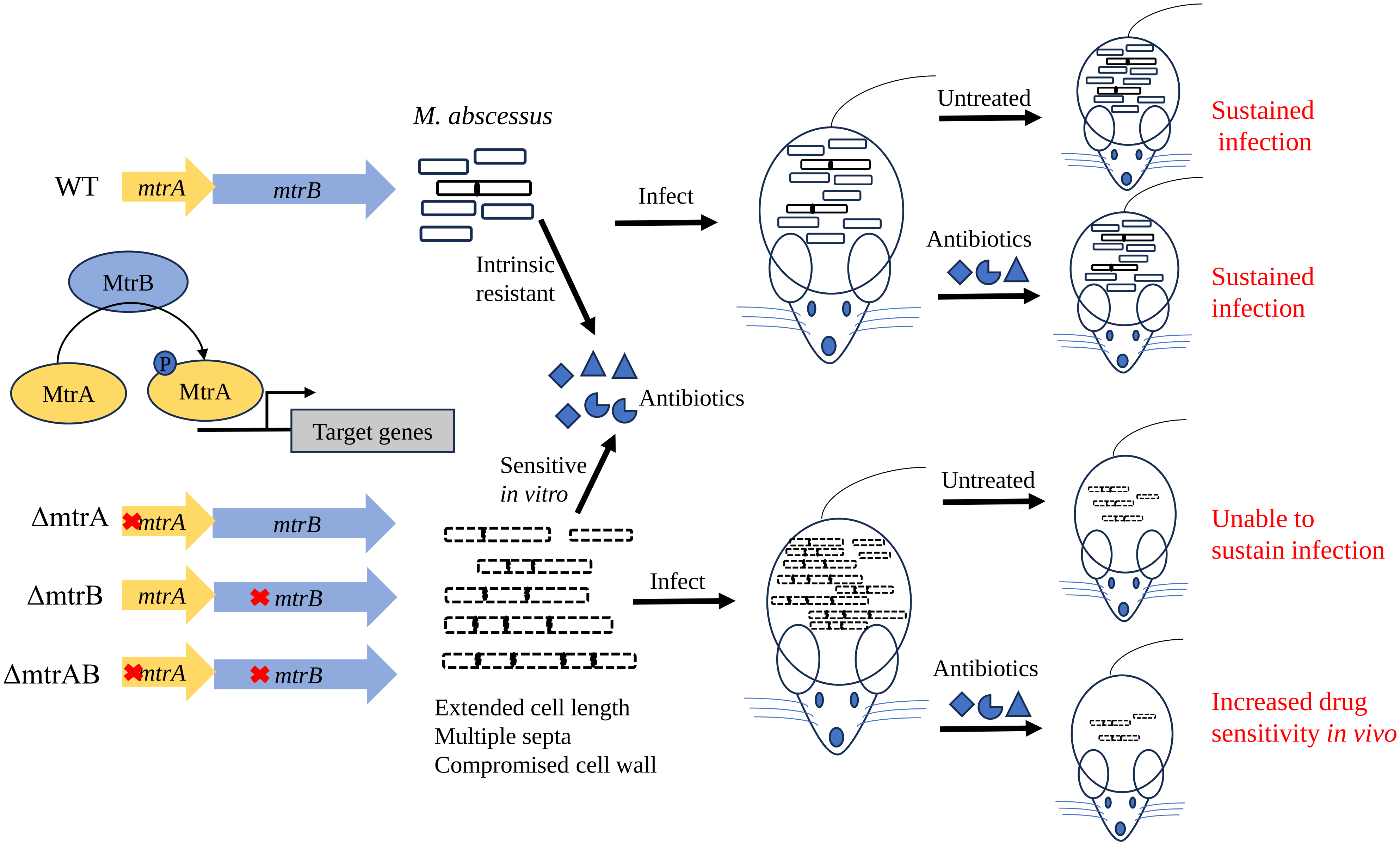
